# Learning mechanism of chromatin domain formation with big data

**DOI:** 10.1101/456525

**Authors:** Wen Jun Xie, Bin Zhang

## Abstract

Chromatin modifications play critical roles in gene regulation and encoding cell phenotypic diversity. The molecular mechanism for their establishment and maintenance is not fully understood due to the complexity of chromatin regulatory pathways. Here we took a data-driven approach and parameterized an information-theoretic model to infer mechanism of chromatin domain formation from genome-wide epigenetic modification profiles. The energy landscape of this model reveals many important chromatin domains that span multiple nucleosomes and exhibit distinct combinatorial patterns of histone modifications, including super (stretch) enhancers, broad H3K4me3 promoter domains, heterochromatin, etc. Transition path analysis further demonstrates that enhancer and promoter domains undergo a sequential maturation process along which the regulatory elements grow from short but stable nucleosome segments to long and potent ones that are modified with many activation marks. On the other hand, the formation of heterochromatin domains is a highly cooperative process, and no intermediate states were found along the transition path. Interaction energies of the information-theoretic model further suggest that heterochromatin domains adopt collapsed, globular three-dimensional conformations that can be stabilized by phase-separated liquid droplets.Our results demonstrate the usefulness of statistical mechanical models and molecular biophysical approaches in interpreting the rich information encoded in epigenomics data.

## Introduction

A remarkable achievement of multicellular organisms is the formation of distinct cell types from the same copy of DNA sequence. Epigenome, the vast number of post-translational modifications to histone proteins that package the DNA into nucleosomes, is expected to play a key role in encoding the cell type diversity.^1,2^ Histone modifications are cell-type-specific and are known to significantly impact gene expression by modulating chromatin structure and recruiting chromatin remodeling complexes.^3^ The importance of these modifications has led to the *histone code* hypothesis, ^4,5^ which states that unique combinations of histone modifications encode distinctive biological outcomes. We present a statistical mechanical model parameterized with epigenomics data to systematically identify histone modification patterns, to quantify their stability against random fluctuations, and to investigate the underlying mechanism for their formation.

Large-scale sequencing studies have provided valuable data on the genome-wide distribution of epigenetic marks.^6^ Numerous approaches have been developed to analyze these data and to extract unique patterns of histone modifications.^^7–9^ Focusing on the correlation between epigenetic marks within the same nucleosome, these approaches have discovered many novel chromatin features, also known as chromatin states. However, it is often the case that epigenetic marks can spread across multiple nucleosomes to form chromatin domains, such as super (stretch) enhancers, 10,11^ broad H3K4me3 domains, ^12^ and heterochromatin.^13^ It is, therefore, crucial to take into account inter-nucleosome correlations to study histone modification patterns beyond single nucleosomes.^14^

The functional importance of histone modifications necessitates the presence of robust mechanisms for their establishment and maintenance. ^15^ Ensuring sustained stability of these chemical modifications within a cell cycle and through cell division can be a challenging task, however. Unlike the DNA sequence, histone modifications are intrinsically dynamic and face constant perturbations from enzymes that add or remove these marks.^16^ Furthermore, roughly half of the histone proteins will be replaced with fresh copies that are arbitrarily modified as the DNA replicates, and these random additions can dramatically alter the epigenome of the daughter cells if left uncorrected. ^^17–19^ Though significant insights on the robustness of histone modifications can be gained by studying chromatin regulatory pathways, 13,20^ a complete understanding based on these pathways is lacking since many of the molecular players that are crucial for epigenome stability and inheritance remain unknown.

We propose an information-theoretic model to identify chromatin domains with distinct histone modification patterns and infer the mechanism for their formation from epigenomics data. This model can be derived and parameterized following the maximum-entropy principle. It rigorously accounts for both intra- and inter-nucleosome correlation between epigenetic marks, and succeeds in identifying well-known chromatin domains, including promoters, enhancers, transcribed genes, heterochromatin, etc. Detailed analysis of these domains revealed that promoters and enhancers could be stabilized at a continuous range of genomic lengths, and more extended domains often grow from shorter ones through a sequential maturation process. On the other hand, heterochromatin domains exhibit a bistable behavior, and the nucleosomes undergo an all or none cooperative transition as they become methylated without experiencing intermediate states. By correlating the model interaction energies with contact probability between nucleosomes in the three-dimensional (3D) space, we found that our model supports condensed, globular heterochromatin conformations that resemble phase separated liquid droplets.

## Results

### Information-theoretic model predicts long-range inter-nucleosome correlations

Chromatin immunoprecipitation followed by deep sequencing (ChIP-Seq) is a powerful method for characterizing the epigenome at high resolution, and has helped to determine genomewide profiles of numerous epigenetic marks across hundreds of cell types.^6,21^ Here, we introduce an information-theoretic approach to analyze these data and investigate the interrelationship of epigenetic marks.

The information-theoretic model describes a chain of *N* interacting nucleosomes, each one of which is characterized by a total of 12 epigenetic marks (Fig. 1). These marks are selected for a comprehensive characterization of the chromatin landscape, and their biological importance are explained in the supporting information (SI). The model’s potential energy adopts the following form
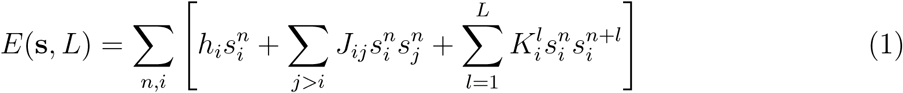

**Figure 1:**
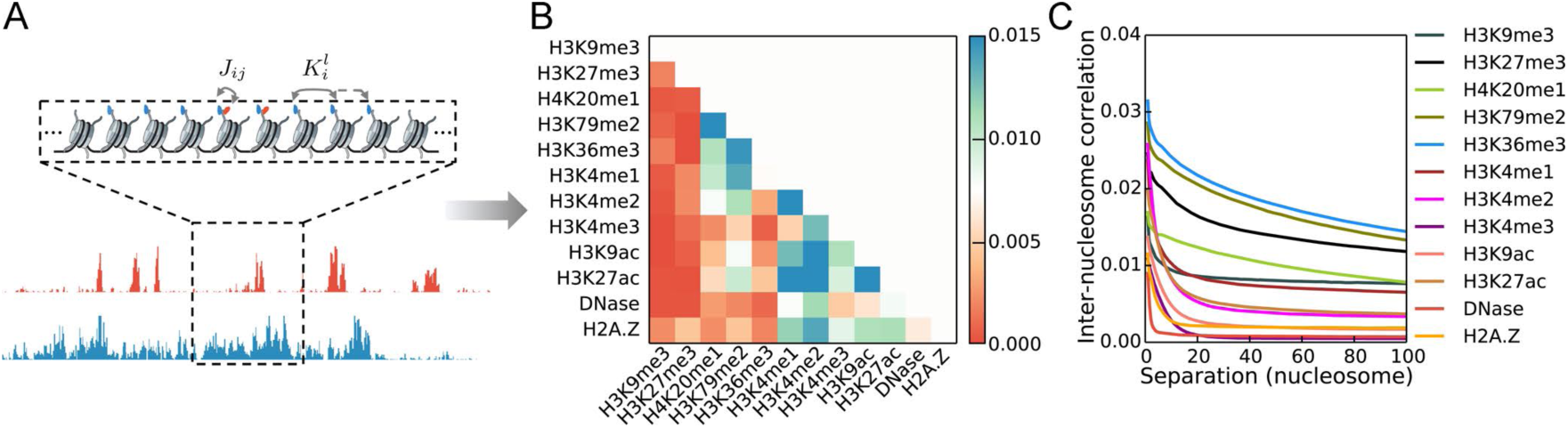
Epigenetic marks exhibit strong intra- and inter-nucleosome correlations. (A) Illustration of the information-theoretic model for studying chromatin domain formation (top panel). The model explicitly considers intra-(*J_ij_*)and inter-nucleosome 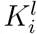 coupling between epigenetic marks, and the values of the coupling strengths are derived from epigenomics data (bottom panel). The fundamental unit of this model is a 200-bp-long genomic segment that includes both the core nucleosome and the linker DNA. A total of 12 epigenetic marks is used to describe the state of each segment. For simplicity, only two marks are shown. (B) Pair-wise correlation between epigenetic marks on the same nucleosome. (C) Self-correlation for epigenetic marks on different nucleosomes as a function of the genomic separation.

The binary variable 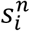 indicates whether the *i*-th mark is present (= 1) at the *n*-th nucleosome or not (= 0). The parameters *h_i_*, *J_ij_* and 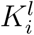 measure the overall propensity for the appearance of the *i*-th mark, the coupling strength between marks *i* and *j* on the same nucleosome, and the coupling strength between the same mark *i* separated by *l* nucleosomes, respectively. As shown in the SI, *E*(s, *L*) is the most probable model that maximizes the information entropy while reproducing the experimental mean and pair-wise correlation of epigenetic marks. Maximum entropy models have been successfully applied to study a wide variety of problems, including protein structure prediction and genome folding. ^^22–28^ A closely related model with only intra-nucleosome correlations has been proposed by Zhou and Troyanskaya to study chromatin states. 9,29^ The added inter-nucleosome interactions here are crucial for studying the spreading of epigenetic marks and the formation of chromatin domains.

As detailed in the *Section: Materials and Methods,* parameters in *E*(s, *L*) can be derived using a Boltzmann learning algorithm. ^30^ For efficient computational sampling, we limited the system size *N* to 25 nucleosomes and adopted the periodic boundary condition. As shown in Table S1, the correlation length for most epigenetic marks is significantly less than 25. In the meantime, the periodic boundary condition ensures that the long-range correlation between some epigenetic marks that give rise to large, periodic chromatin domains can be modeled accurately in a finite system.

To probe the effect of inter-nucleosome interactions and identify the minimalist model with the least parameters that succeeds in capturing the correlation between epigenetic marks, we studied a series of systems with increasing *L.* Figs. 2A and 2B present the results obtained from a model without any inter-nucleosome interactions (*L* = 0). As in-dicated by the blue dots, this model succeeds in reproducing mean mark occupancy and intra-nucleosome correlations, both of which are provided as experimental constraints. The relative error, which is defined as the total absolute difference between experimental constraints and modeled values normalized by the sum of experimental constraints, is less than 5%. Furthermore, higher-order intra-nucleosome correlations, which are measured by the population of all the possible combinatorial patterns formed by the 12 epigenetic marks and are not included as experimental constraints, are accurately predicted as well (Fig. 2A green). On the other hand, the simulated inter-nucleosome correlations between all pairs of epigenetic marks separated by less than 13 nucleosomes differ significantly from experimental values (Fig. 2B).

**Figure 2:**
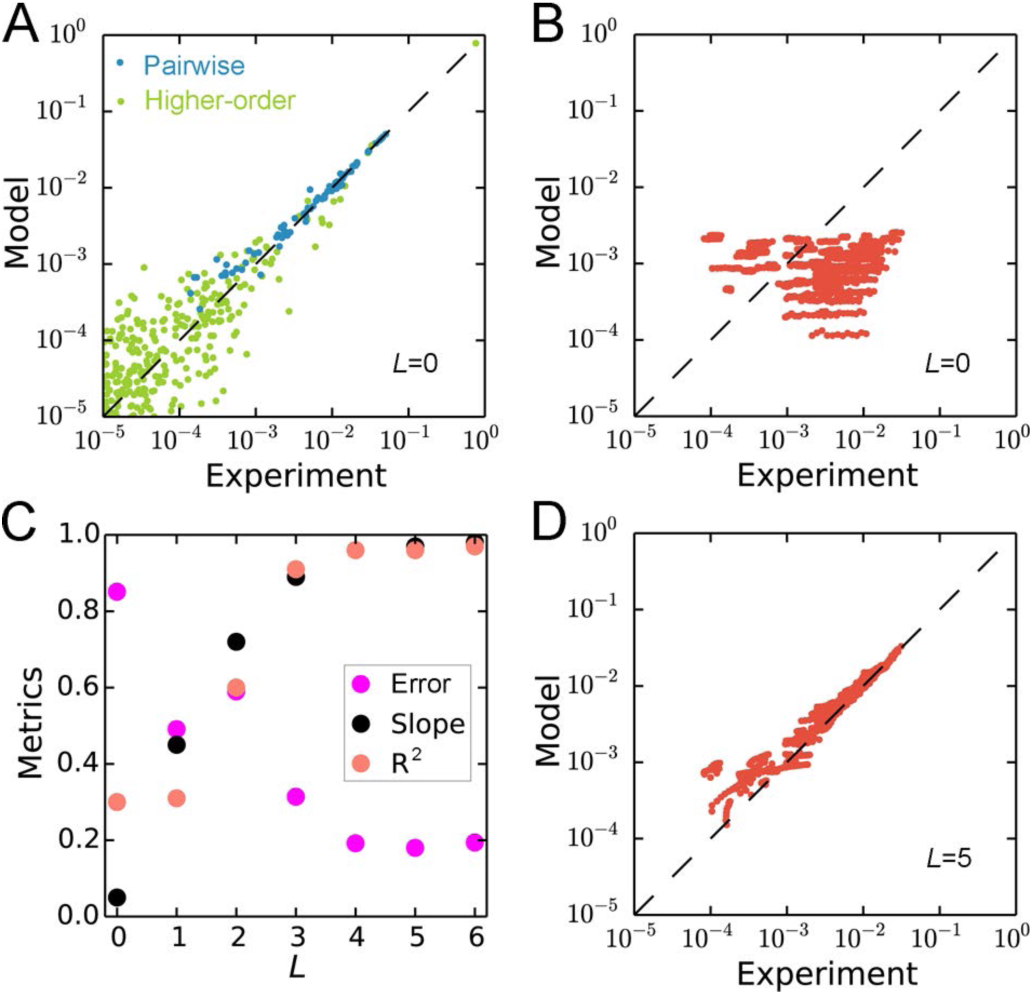
Parameterization and validation of information-theoretic models with different inter-nucleosome interaction cutoff length *L.* (A) Comparison between experimental and simulated (*L* = 0) pair-wise (blue) and higher-order intra-nucleosome correlations (green). (B) Comparison between experimental and simulated (*L* = 0) inter-nucleosome correlations for all pairs of epigenetic marks separated by less than 13 nucleosomes. (C) Quantitative measurements of the performance for models with different *L.* (D) Comparison between experimental and simulated (*L* = 5) inter-nucleosome correlations. The same experimental data as in part B are used for plots in C and D.

The performance of models that explicitly consider inter-nucleosome coupling are shown in Figs. 2C and 2D. We measured the model performance using the quality of the linear regression between simulated and experimental inter-nucleosome correlations, and the relative error between the two. As shown in Fig. 2C, the slope and R-squared of the linear fit improves systematically, and the difference between simulation and experiment continuously decreases up to *L* = 5. Explicit results for the *L* = 5 model are provide in Fig. 2D. Further increasing *L* or introducing inter-nucleosome cross-mark coupling to the energy function *E*(s, *L*) (see Fig. S2) does not improve the model’s performance. We therefore conducted all following analyses using the model with *L* = 5.

### Information-theoretic energy landscape supports chromatin domain formation

A central goal for computational analysis of epigenomics data is to systematically characterize combinatorial patterns of histone modifications that are of functional importance and appear more frequently than random. Previous studies have focused on histone modifications within the same nucleosome to identify chromatin states.^8,29^ Here, we investigate whether distinct patterns also emerge across multiple nucleosomes to form chromatin domains. Towards that end, we searched for basins of attractions supported by the energy landscape of the parameterized information-theoretic model. These energy basins offer representative arrangements of epigenetic marks that are of high population and are ideal candidates for chromatin domains. In the following, we will focus our discussion on the top 100 most populated chromatin domains identified with the steepest descent algorithm (see SI), since they cover over 94% of the whole genome.

Fig. 3A illustrates the average epigenetic mark profiles for the top 100 chromatin domains. The most common domain, which we term as the ground state, exhibits no histone modifications. Heterochromatin domains are identified with signature methylation marks H3K9me3 or H3K27me3. Chromatin domains with H3K4me1 or H3K4me3 are assigned as enhancer and promoter, respectively, though they often share additional activation marks that include H3K4me2, H3K9ac, H3K27ac, DNase, and H2A.Z. A distinct set of chromatin domains (promoter/enhancer) with both H3K4me3 and H3K4me1 marks are found as well, supporting the fundamental similarity between promoters and enhancers. ^31^ Domains marked with H3K36me3, H3K79me2 and H4K20me1 modifications are transcribed gene regions. We further label domains that consist of both gene silencing marks (H3K27me3) and activation marks H3K4me1/2/3 as bivalent, domains marked with transcribed gene marks and regulatory marks as intragenic regulator, and domains marked with H3K9me3 and H3K36me3 as zinc finger protein (ZNF) gene.^6^

**Figure 3:**
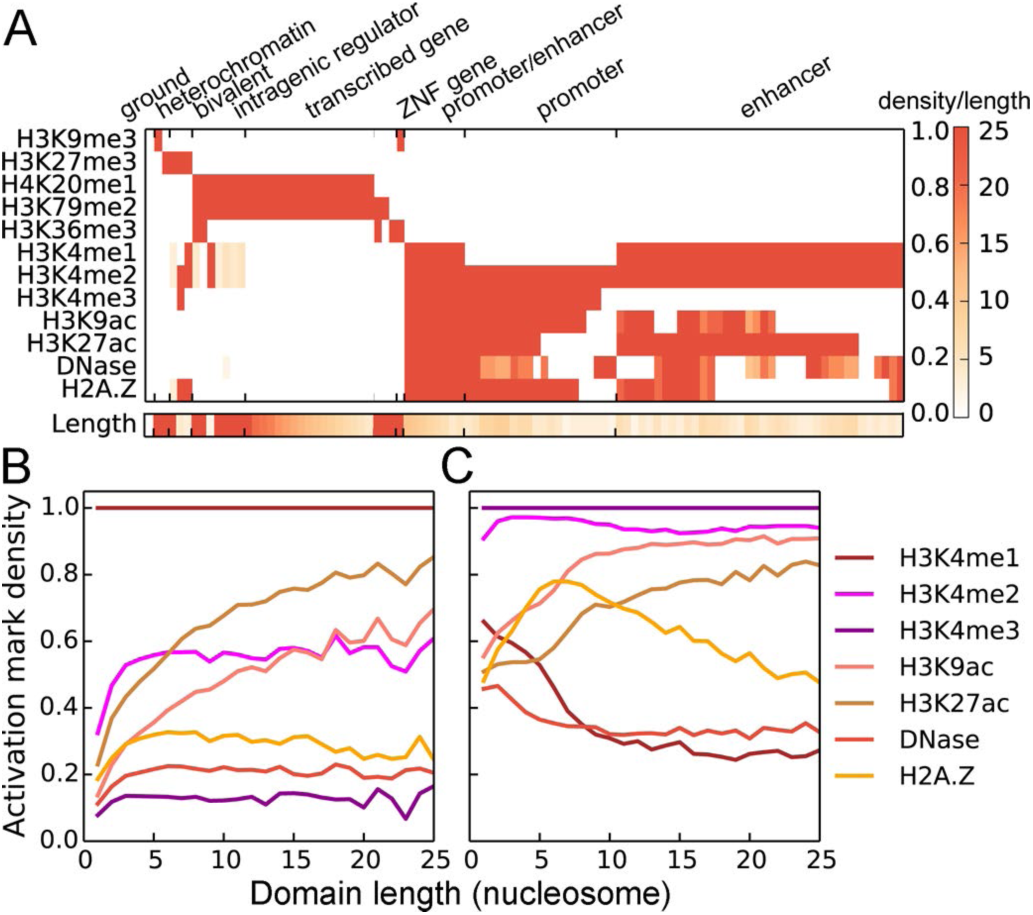
Characterization of chromatin domains predicted by the information-theoretic model *E*(s, *L* = 5). (A) Average epigenetic profiles over modified nucleosomes for the top 100 most populated chromatin domains. The number of modified nucleosomes for each domain is shown in the bottom panel, and the color bar is shown on the side. (B, C) Average probability of observing different epigenetic marks as a function of the length of chromatin stretches with continuous H3K4me1 marks and H3K4m3 marks determined from ChIP-Seq data.

In addition to their unique histone modification profiles, chromatin domains exhibit distinct length dependence as well, as shown in the bottom plot of Fig. 3A. The length of a chromatin domain is defined as the number of nucleosomes with at least one epigenetic mark. For several heterochromatin and transcribed gene domains, all the 25 nucleosomes share the same set of epigenetic modifications. These periodic patterns are consistent with the formation of large silent or transcribed genomic regions that span tens of kilobase pairs. Their periodicity, however, also suggests that these chromatin domains have the potential to spread over the entire genome, and additional mechanisms beyond epigenetic mark interactions, such as boundary elements, must be in play to confine them at specific genomic regions.^32^

On the other hand, we find a series of aperiodic promoter and enhancer domains of varying lengths. We note that extended regulatory domains have indeed been observed previously,^10–12^ and are often termed as broad H3K4me3 domains and super (stretch) enhancers. The increased appearance of additional activation marks in more extended domains (see Figs. 3B and 3C) may support open chromatin conformations to enhance the transcription consistency or the overall expression level of their target genes.

As chromatin domains are local energy minima, their histone modification patterns and length dependence can be understood from the underlying energy landscape. In Fig. 4, we plot the intra- and inter-nucleosome interaction energies between epigenetic marks. Consistent with the complex promoter and enhancer domain patterns, most of the interaction energies between activation marks are negative (blue), supporting their co-existence within the same nucleosome. A notable exception is a strong repulsion between H3K4me1 and H3K4me3, two mutually exclusive marks for defining enhancers and promoters respectively.^6^ We emphasize that the intra-nucleosome energies probe direct interactions among epigenetic marks that correspond better with physical interactions between histone modification enzymes. The correlation coefficients presented in Fig. 1, on the other hand, could arise from a transit effect as a result of indirect couplings. ^29,33^

**Figure 4:**
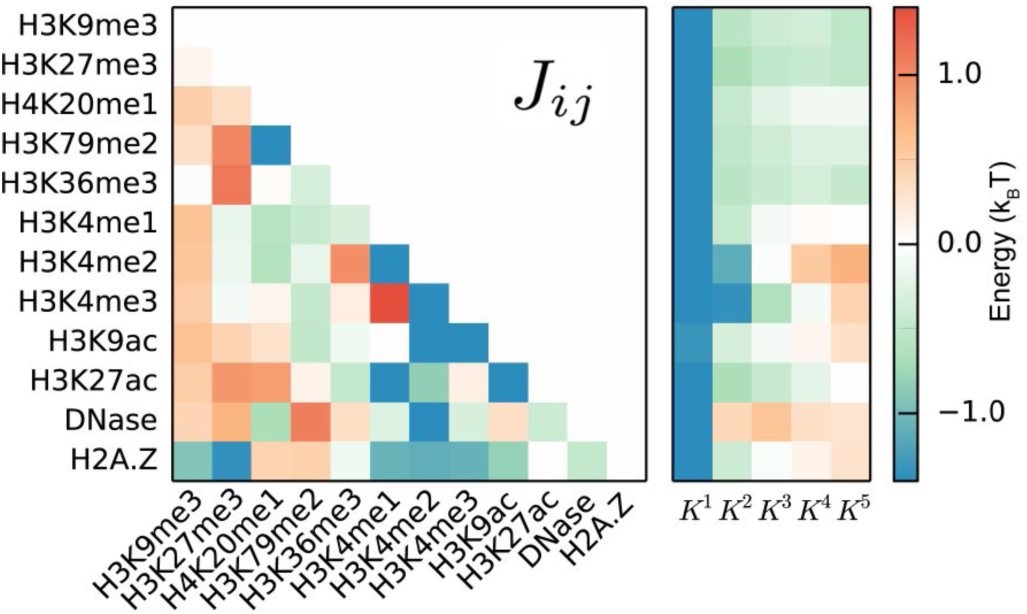
Intra- (Left) and inter- (Right) nucleosome coupling energies between epigenetic marks for the information-theoretic model *E*(s, *L* = 5).

As shown in the right panel of Fig. 4, all epigenetic modifications exhibit attractive interactions between the nearest neighbor nucleosomes (*K*^1^). This attractive interaction decays rather quickly for active marks, explaining the small length of promoter and enhancer domains. On the other hand, the interaction between epigenetic marks for transcribed gene regions or heterochromatin persists over an extended range to promote the formation of large domains.

### Mechanism of chromatin domain formation from transition path calculations

As basins of attraction, chromatin domains are inherently stable and able to withstand transient fluctuations caused by addition and removal enzymes within a cell cycle. Thus, once established, chromatin domains provide a robust mechanism for regulating gene expression and maintaining genome stability. However, the molecular mechanism for their *de novo* creation as cells differentiate and their reestablishment when cells divide remains elusive. We here perform kinetic analysis of the information-theoretic model to provide mechanistic insight into chromatin domain formation.

Towards that end, we first built a kinetic network to explore the dynamical transition between chromatin domains (see Fig. 5A). Each node in this network corresponds to one of the top 100 most populated chromatin domains. A connection between two nodes is introduced if direct transitions between them were observed in a long-time enhanced simulation conducted with the generalized Wang-Landau algorithm.^34^ Transition rates between chromatin domains were further estimated using transition states identified in the simulation trajectory with the transition state theory. ^35^ The detailed algorithm for rate calculations is provided in the *Section: Materials and Methods.* For simplicity, only transitions with energetic barriers less than 15 *k_B_T* are shown in Fig. 5A. A notable feature of this network is its apparent modularity, and chromatin domains with similar biological functions naturally organize into highly connected clusters. The slow transition between clusters highlights the robustness of the overall organization of the epigenome in a given cell type to ensure stable gene expression.

**Figure 5:**
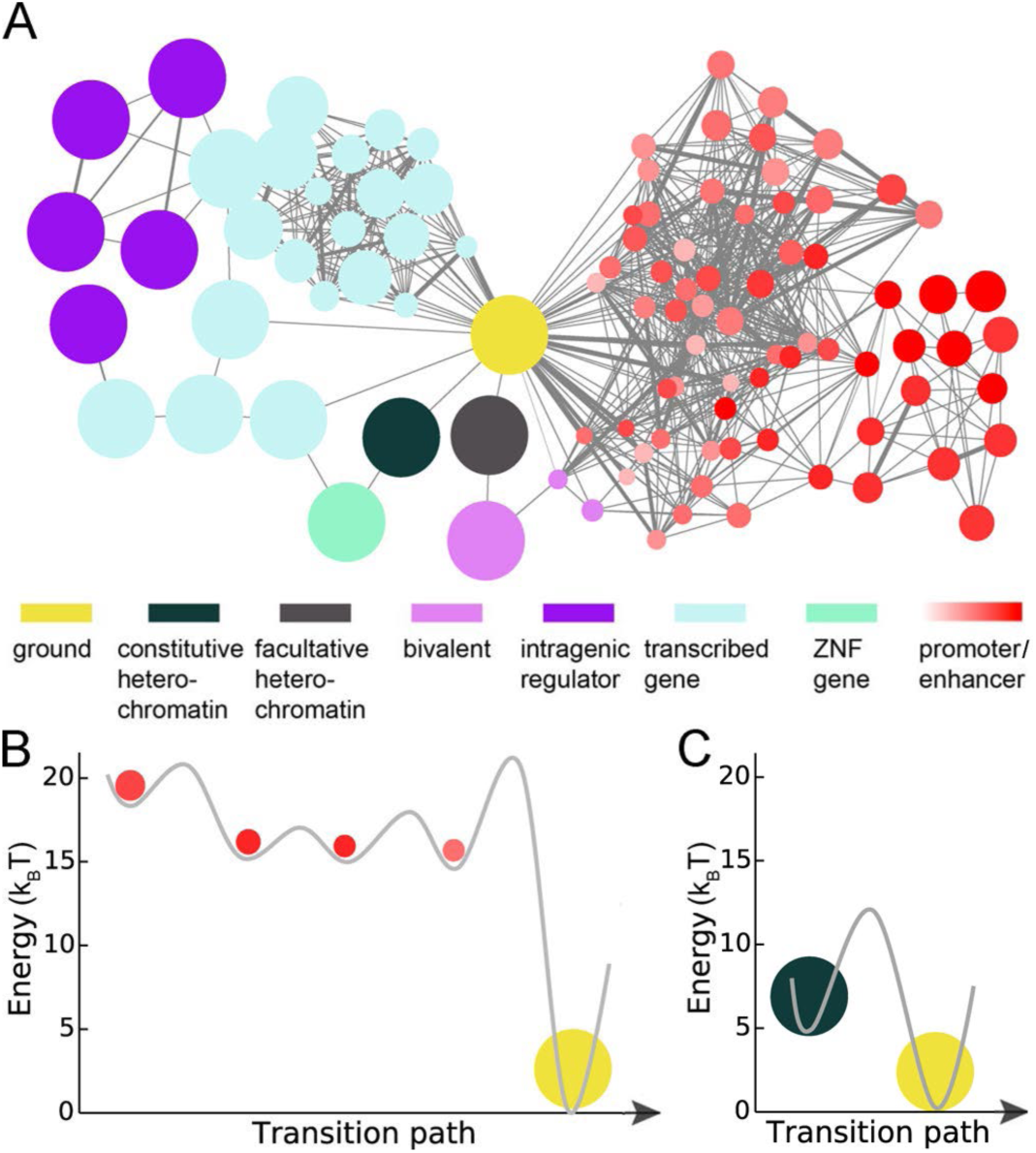
Kinetic analysis of the information-theoretic model *E*(s, *L* = 5). (A) Illustration of the kinetic network for chromatin domain transitions. Each node represents a chromatin domain, and the size of the node is proportional to the domain length. The edge width indicates the value of the transition rates between the two connecting nodes, with thicker lines for faster transitions. The color gradient for promoter and enhancer domains is used to illustrate the densities of activation marks. (B) Energy profile of the most probable transition path from the ground state to an enhancer domain. The line connecting the minima and transition states is provided as a guide for the eye. The length and density of activation marks at each minimum are indicated by the node size and color gradient as in part (A). (C) The most probable transition path from the ground state to the H3K9me3 heterochromatin.

Assuming Markovian dynamics, the time evolution of this kinetic network can be solved analytically to study the transition between any pairs of chromatin domains without performing computationally expensive long timescale simulations.^36^ We expect the Markovian assumption to hold well given that the network nodes are local minima where the system will reside for a significant period to lose memory of the past and achieve dynamical decoupling.

To study the molecular mechanism for the formation of stretch enhancers, we solved the kinetic network and determined the most probable transition path from the ground state to an enhancer domain that extends over 6 nucleosomes. Details of the transition path calculations are provided in the SI. As shown in Figs. 5B and S3, we observe a sequential transition along which the chromatin becomes more open and more enriched with activation marks while the length of the domain grows. Such a multi-step, gradual transition indicates that stretch enhancers may not directly emerge from a genomic region along evolution or as cells differentiate, but instead will undergo a maturation process. The presence of enhancer domains with different potency could help cells fine-tune gene expression levels at various developmental stages. Furthermore, since the backward transition path from the enhancer domain to the ground state is identical to the forward one, the sequential transition suggests that stretch enhancers are more stable with respect to perturbations and are less likely to transition into the ground state and disappear from the chromatin landscape. The increased stability is in accord with their functional importance. Similar conclusions can be drawn from the transition path for the formation of a promoter domain (see Fig. S4)

Next, we determined the transition path from the ground state to the periodic heterochromatin domain with H3K9me3 marks. As shown in Fig. 5C, we find that heterochromatin formation is a cooperative process, and there are no intermediate states along the pathway. To ensure the robustness of this observation, we further determined the transition path using a path deformation algorithm ^37^ that can search for the entire phase space and is not limited by the 100 domains included in the kinetic network. As shown in Fig. S5, the new path shares the same transition barrier and does not exhibit any intermediate state either. The analysis for the H3K27me3 heterochromatin domain supports similar conclusions (see Fig. S6). Cooperativity between nucleosomes will give rise to a bistable system, in which either all or none of the nucleosomes will become methylated, a phenomenon that has indeed been observed in many regulatory networks proposed for heterochromatin formation. ^38,39^ Collective behavior between nucleosomes will significantly enhance the stability of the heterochromatin to ensure a robust inheritance of methylation marks across cell cycles. We will explore possible molecular mechanisms for such cooperativity in the next section.

### Inter-nucleosome interactions support condensed, globular heterochromatin conformations

A striking finding from Fig. 4 is the presence of strong, attractive inter-nucleosome interactions. These interactions measure the propensity for a pair of nucleosomes to share the same epigenetic mark, and can arise if the two nucleosomes are in 3D contact to promote the transfer of corresponding modification enzymes ^38,40^ or enzyme-enzyme interactions ^39^ (see (Fig. 6A). Consistent with this interpretation, all epigenetic marks exhibit a strong *K*^1^ interaction energy as a result of the spatial proximity between the two neighboring-in-sequence nucleosomes. Furthermore, the interaction energies typically decay at larger genomic separations as the nucleosomes become farther apart in space. It is therefore tempting to assume that internucleosome interaction energies are proportional to the contact probability between pairs of nucleosomes. Since different polymer configurations exhibit distinct trends in the variation of contact probabilities as a function of sequence length, studying the scaling behavior of interaction energies offers the opportunity to derive 3D chromatin domain conformations. Indeed, we find that interaction energies of the heterochromatin mark H3K9me3 decay slower than that of the activation mark H3K4me3, consistent with the fact that heterochromatin domains are more condensed than euchromatin.

**Figure 6:**
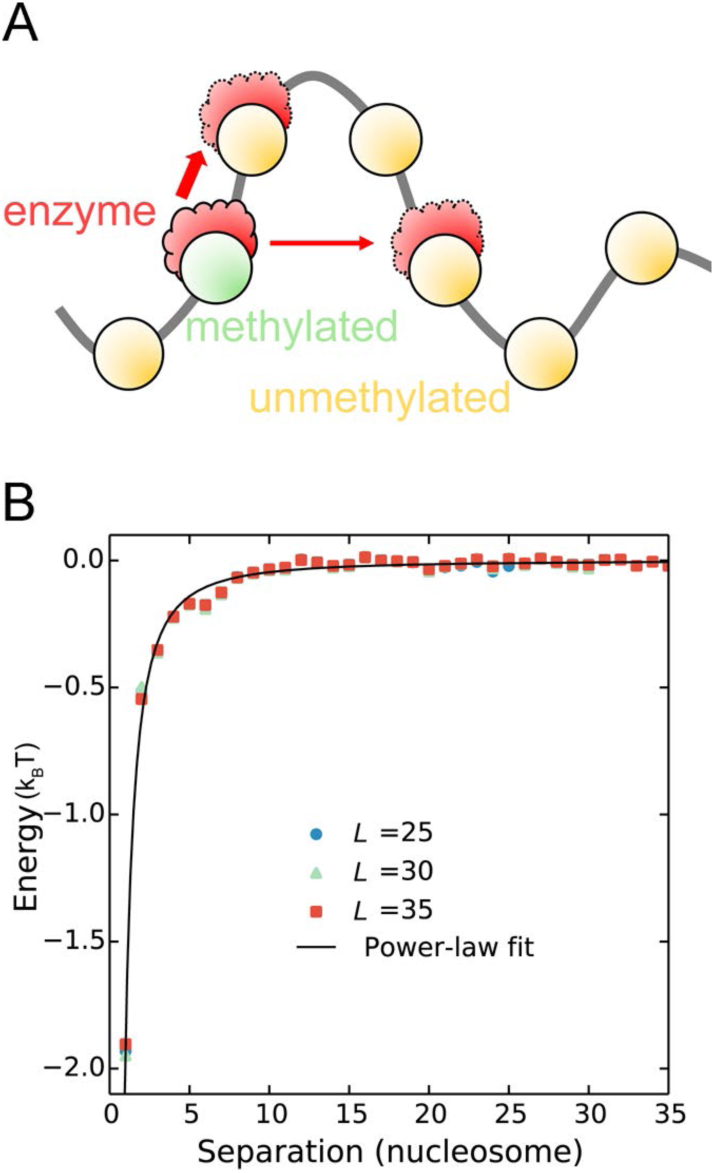
Inferring heterochromatin formation mechanism from inter-nucleosome interaction energy. (A) Illustration of the transfer of histone-modifying enzymes (red) from the methylated nucleosome (green) to others that are close in 3D space. This enzyme transfer is promoted by diffusion and its rate is impacted by the distance between nucleosomes that increases at larger sequence separations. (B) Inter-nucleosome interaction energy for epigenetic mark H3K9me3 as a function of the genomic distance.

Quantitatively inferring scaling exponents from Fig. 4 can be challenging, however, due to the limited number of inter-nucleosome couplings included. To more accurately determine the value of interaction energies at large genomic separations, we studied an additional model consisting of 201 nucleosomes. We only included the H3K9me3 mark for computational efficiency, and each nucleosome can either be methylated or free of modifications. Since H3K9me3 does not co-localize strongly with other epigenetic marks, this single mark model is expected to provide a good description for heterochromatin formation.

The single mark model adopts the same Hamiltonian as in Eq. 1. We included internucleosome interactions to a much longer range than the multi-mark model, and determined the parameters again using the Boltzmann learning algorithm. Fig. 6 presents the resulting interaction energies as a function of nucleosome separation. These energies are robust and insensitive to the cutoff length *L*, as well as the parameters used to process and binarize the raw data (see Fig. S7). We further fitted the interaction energies for *L* = 35 with a power law expression *E*(*l*) = *al^α^*, and obtained a value for *α* = –1.61(±0.07). An exponent that falls in the range between -2 and -1 has been shown to support a phase transition in the one-dimensional Ising model,^41^ giving rise to the bistability seen in Fig. 5C. Assuming that interaction energies are proportional to the contact probability *P*(*l*), an exponent of -1.5 suggests that the heterochromatin conformation is consistent with an equilibrium globule. ^42^ Globular conformations differ significantly from the rigid fibril structures with a diameter of 30 nm, and can be stabilized by phase-separated liquid droplets formed by heterochromatin protein 1 (HP1).^43–47^

## Discussions

Genome-wide histone modification profiles provide a comprehensive characterization of the chromatin landscape. Using an information-theoretic model and rigorous statistical mechanical tools, we demonstrated that epigenomics data could shed light on the mechanism of chromatin domain formation as well. In particular, we found that the H3K9me3 heterochromatin domain exhibits bistability and forms in a highly cooperative process. Furthermore, the 3D conformation of heterochromatin is consistent with a globular conformation, but not the rigid fibril structures. We note that determining chromatin structure at the kilo-base range is extremely difficult, and the existence of 30 nm chromatin fibers *in vivo* remains controversial,^48–50^ though increasing evidence argues against their presence.^51–53^ We contribute to resolving this mystery indirectly by studying the role of chromatin conformation in chromatin domain formation.

Our study supports an essential role of chromatin folding in facilitating the spreading of epigenetic marks across nucleosomes. Explicit consideration of chromatin conformational dynamics and its coupling with the chemical reaction dynamics of histone modifications will be critical for a faithful description of heterochromatin formation that drives simultaneous transitions in both the conformational and chemical space.

## Methods

### Parameterization of the information-theoretic model

We obtained genome-wide histone modification profiles for the IMR90 cell line from the ROADMAP epigenomics project.^6^ These data were binarized with a Poisson background model at a resolution of 200 bp that corresponds to the nucleosome repeat length. From the binarized data, the mean occupancy for epigenetic mark 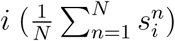, the intra-nucleosome mark correlations 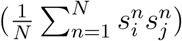, and the correlation coefficient between marks *i* and *j* that are separated by *l* nucleosomes 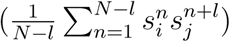 were calculated, where *N* is the genome length.

Using the mean occupancy and mark correlations as experimental constraints, parameters in the energy function *E*(s, *L*) defined in Eq. 1 can be determined with a Boltzmann learning algorithm.^30^ This algorithm iteratively updates the parameters using the difference between the experimental correlation coefficients and simulated ones estimated using replica exchange Markov chain Monte Carlo sampling. ^54^ The iteration terminates when the relative error 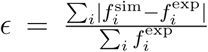 is less than 5%, where *f_i_* include mean epigenetic mark occupancy and pair-wise correlation coefficients. A detailed description of the parameterization algorithm is provided in the SI.

### Calculation of transition rates between chromatin domains

We used Monte Carlo simulations to probe connections between chromatin domains and to calculate transition rates. Specifically, for a pair of chromatin domains *A* and *B*, we first selected a large set of paths that originated from *A* and ended up in *B* from a trajectory that has sufficiently traversed the entire energy landscape. From the path ensemble 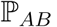, the configuration of the transition state connecting A and B is identified with the energy
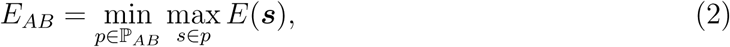

where *E*(***s***) is the energy of configuration ***s*** along the transition path *p.* The energy barrier is then determined as Δ*E_AB_* = *E_AB_ – E_A_* for the transition from *A* to *B,* and Δ*E_BA_* = *E_AB_ – E_B_* for the reverse transition. From these barriers, we estimate the transition rates using the Arrhenius equation 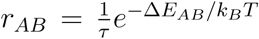, where *τ* is the timescale for observing a fluctuation in histone modification. Noting that it’s non-trivial to efficiently sample the huge phase space. We therefore employed the generalized Wang-Landau algorithm^34^ to bias the simulations for enhanced transitions between domains (see SI for more details).

## Acknowledgement

We thank X. Che for helpful discussions. This work was supported by the National Science Foundation Grant MCB-1715859.

## Author contributions

W.J.X. and B.Z. designed research, performed research, analyzed data, and wrote the manuscript.

## Supplementary Information

## 1. Calculating experimental constraints from genomic profiles of epigenetic marks

For a comprehensive characterization of the chromatin landscape, we analyzed the collective behavior of 12 histone marks. These marks include histone H3 lysine 4 monomethylation/dimethylation/trimethylation (H3K4me1/me2/me3) that are important for identifying enhancer and promoter regions; H3 lysine 9 acetylation (H3K9ac) and H3 lysine 27 acetylation (H3K27ac) that are associated with increased activation of gene promoters and enhancers, respectively; H3 lysine 9 trimethylation (H3K9me3) and H3 lysine 27 trimethylation (H3K27me3) that are signatures of constitutive heterochromatin and facultative heterochromatin; H3 lysine 36 trimethylation (H3K36me3), H3 lysine 79 dimethylation (H3K79me2), and H4 lysine 20 monomethylation (H4K20me1) that are indicative of transcribed gene regions; DNase I hypersensitive sites that probe exposed DNA and open chromatin regions, and an important histone variant H2A.Z.

ChIP-Seq and control data of these epigenetic marks were downloaded from the ROADMAP epigenomics project (1). We focused our analysis for the IMR90 cell line since its data have the highest signal-to-noise ratio percentile. From the raw data, we performed binarization and determined the mean and correlation coefficients of epigenetic marks as detailed in the *Section: Materials and Methods* of the main text.

## 2. Estimating inter-nucleosome correlation length of epigenetic marks

To quantitatively probe the strength of inter-nucleosome interactions, we fitted the correlation coefficients between epigenetic marks as a function of nucleosome separation (*l*) with the expression 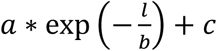. The fitting parameter *b* is used to measure the correlation length of epigenetic marks. Fitting results for self-correlation of the same mark are shown in Table S1.

Most of the correlation lengths are smaller than 25 nucleosomes, supporting our use of *N* = 25 as the system size. For marks with self-correlation lengths longer than 25, they often support the formation of large, homogeneous chromatin domains. The effect of these marks will therefore be well captured by the periodic boundary condition.

## 3. Deriving information-theoretic models from the maximum entropy principle

As mentioned in the main text, *E*(***s***, *L*) is the most probable model to reproduce experimental constraints. Here we provide a detailed derivation for its expression based on the maximum entropy principle.

To start, we define the system of interest as a chain of *N* nucleosomes, each one of which is characterized by *M* epigenetic marks. A configuration, or state, of this system is denoted as 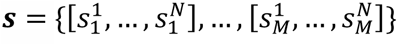 which can be viewed as a *M* × *N* matrix and 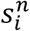 denotes whether mark *i* is present (= 1) at the *n*-th nucleosome or not (= 0). Next, we seek for a probability distribution function of state ***s*** that maximizes the information entropy
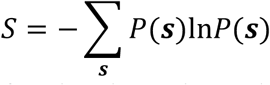

while reproducing the mean occupancy of each epigenetic mark *i*,
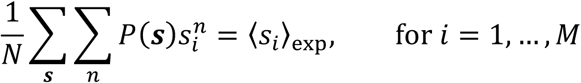

and the correlation between two epigenetic marks *i* and *j* separated by *l* nucleosomes,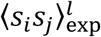,
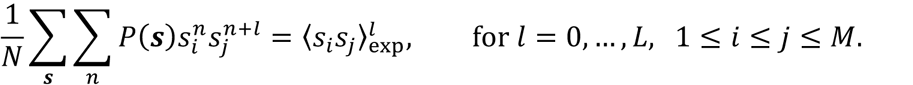

The solution to this constrained optimization problem can be found by minimizing the following Lagrangian
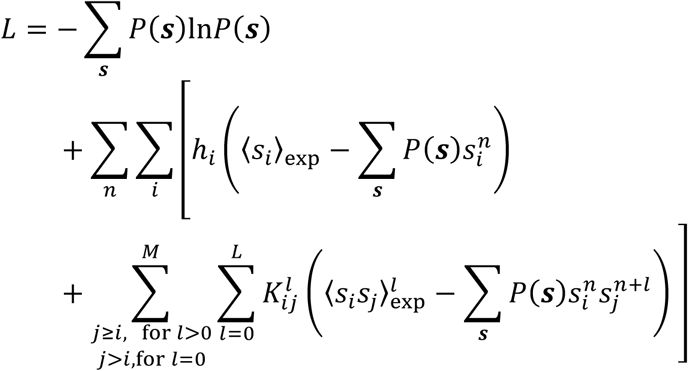

where *h_i_* and 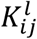 are Lagrangian multipliers.

The solution for 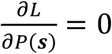 provides the least structured *P*(***s***) in the form of a Boltzmann distribution
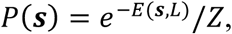

where the energy function is defined as
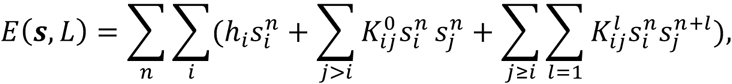

and *Z* is the partition function. The unit of energy is k_B_*T*.

Following the argument above, we can arrive at a series of information theoretic models presented in the main text by adjusting the input experimental constraints. For example, if only 〈*s_i_*〉_exp_ and 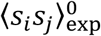, for 1≤ *i < j* ≤ *M* are provided as constraints, then we arrive at the *intra-nucleosome model,*
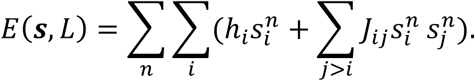

If *L* × *M* additional self-correlation coefficients 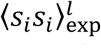, for *i* = 1, ···, *M*, and *l* = 1, ···, *L* are provided as constraints, then we arrive at the *inter-nucleosome model with only self-interactions,*
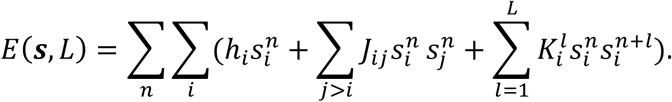

Finally, if we further introduce cross-mark correlation between neighboring nucleosomes 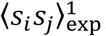, for 1 ≤ *i* < *j* ≤ *M* as constraints, we arrive at the *inter-nucleosome model with both self- and cross-mark interactions,*
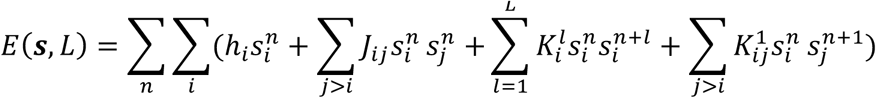

## 4. Parameterizing information-theoretic models with a Boltzmann learning algorithm

Parameters in the maximum entropy model can be derived using a Boltzmann learning algorithm. This algorithm minimizes the cross-entropy
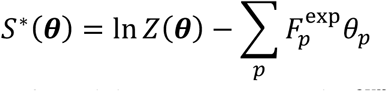

where 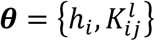 is the collection of model parameters, and 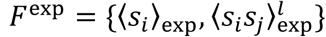 is the collection of experimental constraints. It is straightforward to demonstrate that at the stationary point of *S*^*^(*****θ*****), the simulated ensemble averages match with experimental constraints.

The minimum of *S*^*^(*****θ*****) can be found with the steepest gradient descent method that iteratively updates the parameters with the following expression
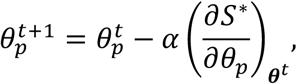

Where the gradient of S^*^ is defined as:
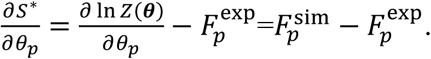

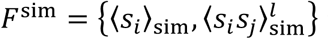 is the collection of simulated ensemble averages for mean mark occupancy and correlation coefficients between marks.

To accelerate steepest gradient descent in the relevant direction and dampen oscillations, a momentum step was included when updating the parameters (2):
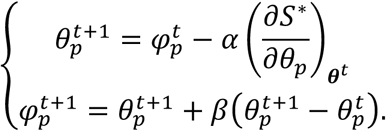

We used a learning rate of *α* = 0.002 and *β* = 0.9. At each iteration *t*, we used the Metropolis Monte Carlo algorithm to sample the Boltzmann distribution and estimate the ensemble averages 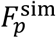. We further applied the parallel tempering technique to enhance sampling efficiency. A total of thirteen replicas was used with temperatures evenly distributed between 0.8 and 2.0, with each replica lasting for 10^6^ Monte Carlo steps per iteration. The parameters were set as zero at the beginning and were updated until the relative error defined as 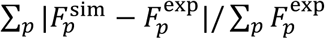 is less than 5%.

## 5. Identifying chromatin domains with the steepest descent algorithm

To identify robust combinatorial patterns of epigenetic marks and chromatin domains, we searched for basins of attractions supported by the information-theoretic model using steepest gradient descent optimization. Like other Hopfield-like systems (3)(4), these basins are local minima that represent patterns with high probabilities of appearance. Local minima are defined as states whose energies are lower than all the neighboring configurations. The neighborhood of a state includes all configurations that differ from the state with only one epigenetic mark. Following this definition, the closest local minimum for any configuration can be found by the steepest gradient descent algorithm. In this algorithm, we iteratively select out the configuration that has the lowest energy in the neighborhood of the lowest energy state from previous iteration until convergence.

To identify the set of local minima that most resemble the configurations observed in genomewide profiles of epigenetic marks, we first partitioned the genome into unmodified and modified regions. The modified regions were identified as stretches of genomic segments that do not contain any gaps longer than 25 nucleosomes. Gap regions do not exhibit any epigenetic marks. The local minimum for unmodified regions will always be the ground state. For modified regions, we further divided them into configurations with 25 nucleosomes that correspond to the system size of our model. For each 25-nucleosome long region, we performed the steepest gradient descent optimization to find the corresponding local minimum (i.e., chromatin domain).

## 6. Estimating transition barriers using Wang-Landau sampling

We used the following approach to determine the transition barriers between pairs of chromatin domains.

*First*, we collected an ensemble of transition paths connecting the two domains from a simulation trajectory conducted using the Metropolis Monte Carlo algorithm coupled with the Wang-Landau algorithm (5). The Wang-Landau algorithm is an enhanced sampling technique that aims at producing a random walk over basins of attraction, thus greatly accelerating the transition between local minima. It estimates the density of states of the system along the simulation and use it to bias the sampling. In the following, we briefly summarize the key steps of this algorithm when applied to a spin glass model.

Suppose that we are interested in the top *N* most populated minima of the spin glass model. By assigning spin configurations to their closest local minimum using the steepest descent algorithm outlined in the previous section, the phase space thus can be partitioned into *N* + 1 basins *B*_0_, *B*_1_, *B*_2_, … , *B_N_*. *B*_0_ includes all the configurations that cannot be assigned to the top *N* minima. To estimate the density of state for each basin, we further partition the energy space *u*_1_ < *u*_2_ < ··· < *u_L_* = ∞ into *L* intervals, where *u*_1_ is the lower bound of the model Hamiltonian, and is set to zero in our case. Therefore, the phase space is now partitioned into (*N* + 1) × *L* regions, *B_im_* = {***s*** ∈ *B_i_*: *E*(***s***) ∈ [*u_m_*, *u*_*m*+1_)}, for *i* = 0, ··· ,*N* and *m* = 1,··· *L* . The statistical weight (density of states), *g_im_*, for each region is set as one initially.

We then update the density of states by performing the following Monte Carlo sampling. Starting from a random configuration s in the basin region *B_im_*, we perform a Monte Carlo move with a single-spin flip. This move will lead to a new configuration ***s***′ in the region *B_jn_*. The new configuration will be accepted with the following probability
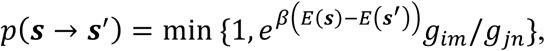

where *β* is an effective temperature.

Along the simulation, we keep track the history of the simulation trajectory with a histogram. Specifically, we will update the number of times a region has been visited *H_jn_*, by 1 if the new configuration ***s***′ is accepted. In the meantime, the corresponding density of states *g_jn_* , will be updated as *g_jn_f* , where *f* is a scaling factor larger than 1. The goal of the Wang-Landau sampling is to achieve a flat histogram, indicating a random walk in the energy basins.

The above sampling will be conducted iteratively. For example, if the flatness of the histogram becomes acceptable (maximal fluctuation is less than 25%.), *f* will be scaled down to 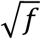. The iteration completes when *f* is close to 1.

When applying the Wang-Landau algorithm to the informational theoretic model, we limited the sampling to the top 100 chromatin domains that cover more than 94% of the nucleosomes. The modification factor *f* of density of states is set as *e* ≈ 2.71828 at the beginning of the sampling. The effective temperature *β* is set as 1/(2k_B_*T*) to accelerate the sampling, resulting the Metropolis acceptance ratio of ~15%. The energy range of interest is [0,120 k_B_*T*] and the energy width is 10 k_B_*T*. We stopped the sampling when *f*<1.000001.

*Second,* from the ensemble of transition paths 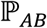 collected from Wang-Landau sampling, the energy for the transition state connecting chromatin domains *A* and *B* is identified as
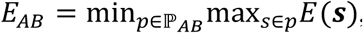

where *E*(***s***) is the energy of configuration ***s***. The transition barrier is then obtained as *ΔE_AB_* =*E_AB_* – *E_A_* for the transition from *A* and *B*, and *ΔE_BA_* = *E_AB_* – *E_B_* for the reverse transition.

## 7. Calculating the most probable path between chromatin domains using the transition path theory

Using the energetic barriers determined from Wang-Landau sampling, we estimated the transition rates between chromatin domains with the Arrhenius equation 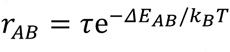, where *τ* is the timescale for a fluctuation in histone modification. From the set of transition rates, we built the kinetic network shown in Fig. 5A. Under the Markovian assumption, this kinetic network fully specifies the long-time dynamics of the system.

To determine the most probable pathway connecting chromatin domains *A* to *B*, we applied the transition path theory to the kinetic network using the software PyEMMA (6). Transition path theory is a probabilistic framework to analyze the mechanism of reaction or transition and provides the reactive rate of state transformation (7). The core concept in transition path theory is the committor probability. The committor probability 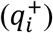 at domain *i* is defined as the probability that the system reaches domain *B* first before visiting domain *A*. Thus, the committor probability can be solved by the simple relation with the transition probability matrix
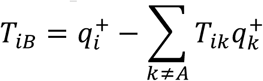

The transition probability matrix ***T*** was constructed from the rate matrix ***r*** using the expression ***T*** = *e*^***r*****Δ*****t***^. A lag time of Δ*t* = 1 was used.

Among all the transition between two domains, only a fraction is reactive. The effective flux of transition between intermediate states *i*→*j* contributing to the path *A*→*B* is
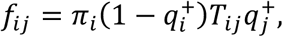

where *π_i_* is the equilibrium distribution of chromatin domain *i* that can be determined from the relation **π*****T*** = **π**. Then we picked up the most probable transition pathway connecting *A* and *B* as the pathway with the greatest flux (*f_ij_*) along it (6).

## 8. Calculating the minimum energy barrier path between chromatin domains using a path deformation approach

As an independent validation of the transition paths determined in the main text, we further determined the transition barrier and minimum energy path between the ground state and the H3K9me3 heterochromatin using a path deformation approach (8)(9). This approach explores the entire phase space when searching for the transition state. Therefore, unlike the method presented in the main text, it is not restricted by the number of chromatin domains included in the kinetic network.

For the convenience of discussion, we define the mark-flipping operator *O* that turns a particular epigenetic mark on or off. Since a continuous path connecting two states *A* and *B* consists of a set of configurations that differ only by one mark, it can be described by a product of operators, *B* = *O*_1_*O*_2_ … *O_n_A*.

The path deformation algorithm starts with a randomly generated minimum length path, for which the number of operators equals to the total number of different mark configurations in the two states. Next, it aims to find the minimum transition barrier connecting the two states by iteratively deforming the highest energy configuration along the path. We used two types of moves to deform the path. The first one swaps the operator that leads to the highest energy configuration with the one leaving it. In the second move, two operators are added before and after the operator that leads to the highest energy configuration. These two operators modify the same mark randomly selected from the 12 epigenetic marks. They have opposite effects: if one switches a mark on, the other will turn it off. Newly deformed paths are only accepted if they lead to a decrease of the transition barrier.

We carried out a total of 1000 independent calculations to search for the transition barrier between the ground state and the heterochromatin domain. For each calculation, the iteration terminates when the total length of the path exceeds 20000 configurations. Remarkably, all these paths give rise to the same barrier as the one presented in the main text (Fig. 5C). From the transition state, we further constructed the minimum energy path using the steepest gradient descent algorithm. As shown in Fig. S5, the path exhibits no intermediate states, supporting a cooperative transition for the formation of constitutive heterochromatin.

## 9. Evaluating the sensitivity of interaction energy strength with respect to the Poisson threshold used in the binarization of experimental data

Genome wide histone modification profiles were processed with a Poisson background model to remove biases that might arise from the DNA sequence (10). A Poisson threshold of 0.0001, a typical number used in the widely popular software ChromHMM (11), was used to detect significant experimental signals compared to the control data. A variation of the threshold value could significantly affect mark occupancy, and the correlation coefficients. Indeed, as shown in Fig. S7A, the inter-nucleosome correlation changes significantly as we vary the threshold by a factor of two.

To investigate the sensitivity of the derived interaction energies with respect to the Poisson threshold, we reparametrized the simplified heterochromatin domain shown in Fig. 6 of the main text using these new correlation coefficients. It is reassuring to see that the inter-nucleosome interaction energies with different thresholds are quite close to each other (see Fig. S7B). Fitting these interaction energies using a power-law expression *E*(*l*) = *al^α^* provides scaling exponents of –1.59 ± 0.06 and –1.45 ± 0.06 for the threshold 0.00005 and 0.0002, respectively. These number are very close to the value –1.61(±0.07) provided in the main text.

---

**Table S1.** Correlation length of epigenetic marks. See text *Section: Estimating internucleosome correlation length of epigenetic marks* for a detailed discussion.

| Epigenetic mark | Correlation length<br>(nucleosome) |
| --- | --- |
| H3K9me3 | 5.6 |
| H3K27me3 | 24.1 |
| H4K20me1 | 53.1 |
| H3K79me2 | 45.1 |
| H3K36me3 | 37.1 |
| H3K4me1 | 5.8 |
| H3K4me2 | 5.9 |
| H3K4me3 | 6.2 |
| H3K9ac | 6.9 |
| H3K27ac | 8.0 |
| DNase | 1.0 |
| H2A.Z | 3.6 |

**Fig. S1.**
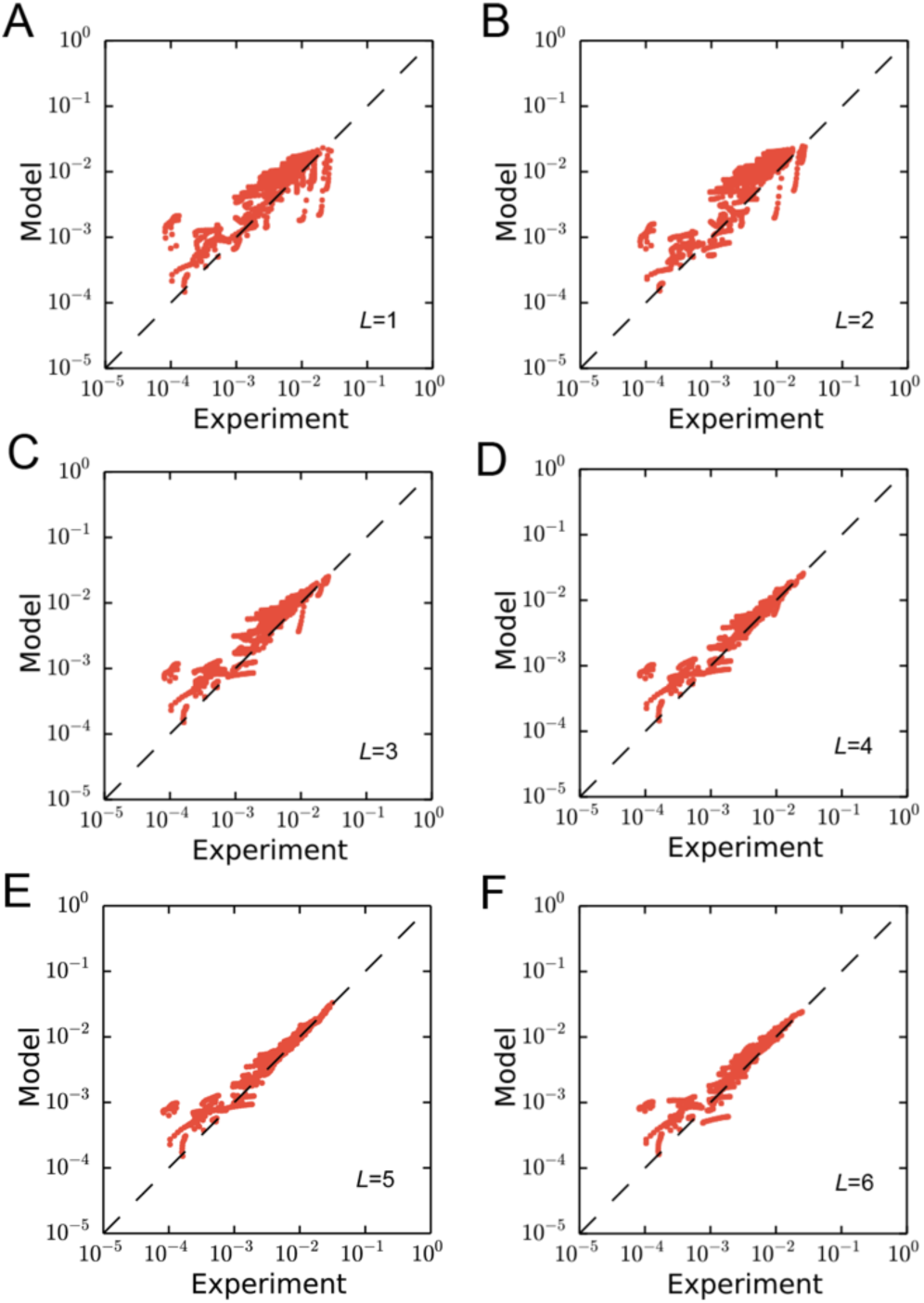
Comparison between experimental inter-nucleosome correlations and predictions from information-theoretic models *E*(*s*, *L*) with different *L*. In making these plots, the same experimental data as in Fig. 2B of the main text was used.

**Fig. S2.**
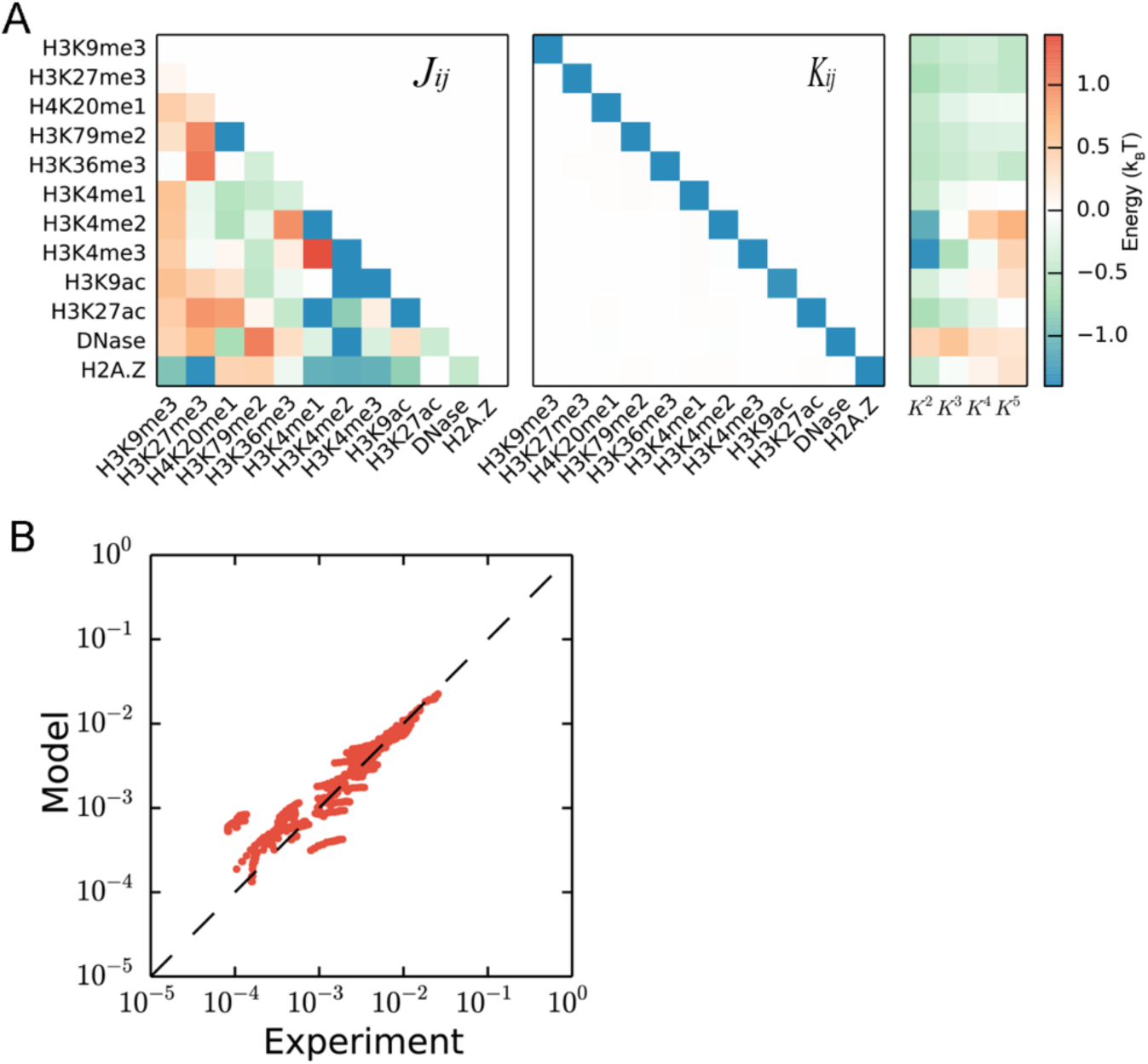
Performance of an information theoretic model with explicit coupling between all pairs of epigenetic marks on neighboring nucleosomes. (A) Intra-nucleosome interaction (*left*), nearest neighbor interaction (*middle*), and long-range interaction (right) energies of the model. An explicit expression of the model Hamiltonian is provided in the *Section: Deriving information-theoretic models from the maximum entropy principle.* The cross-mark interaction energies are significantly smaller compared to the self-interaction, and are barely visible in the middle panel. (B) Comparison between experimental and simulated inter-nucleosome correlations. The experimental data is identical to those shown in Fig. 2B of the main text. The relative error between simulated and experimental data is 17.6%. The slope and R-squared of the linear fit are 0.93 and 0.97, respectively. We therefore conclude that including the cross-mark coupling on the nearest neighbor nucleosomes doesn’t improve the model performance significantly. For simplicity, these terms were not included in the models presented in the main text.

**Fig. S3.**
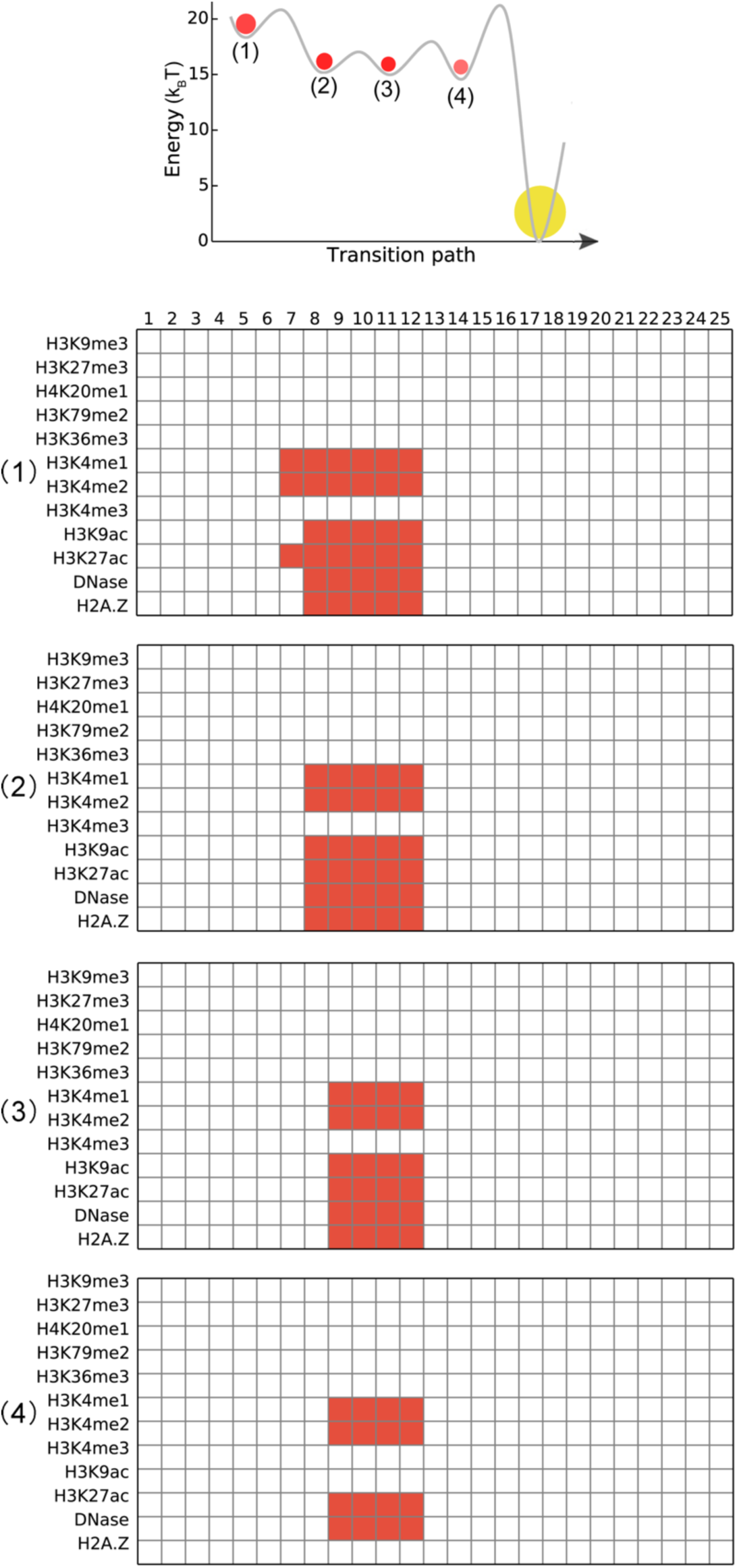
Configurations for intermediate states along the most probable transition path from the ground state to an enhancer domain. The same path as in Fig. 5B is shown at the top for reference. The rows in each configuration correspond to different epigenetic marks, and the columns represent different nucleosomes. A grid is marked with red if the epigenetic mark is turned on for that nucleosome.

**Fig. S4.**
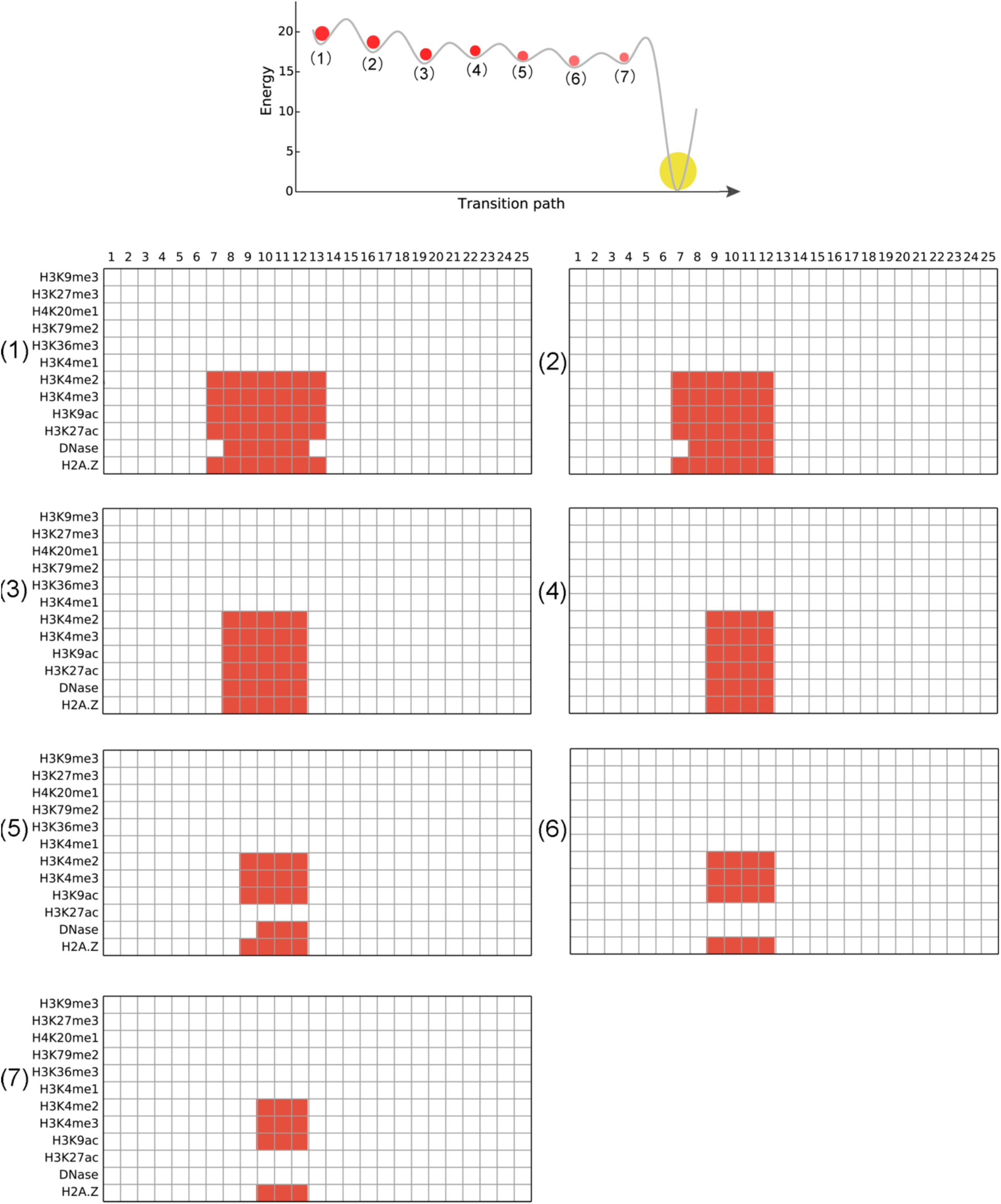
Configurations for intermediate states along the most probable transition path from the ground state to a promoter domain. Similar to the path shown in Fig. 5B of the main text, this path exhibits multiple intermediate states as well.

**Fig. S5.**
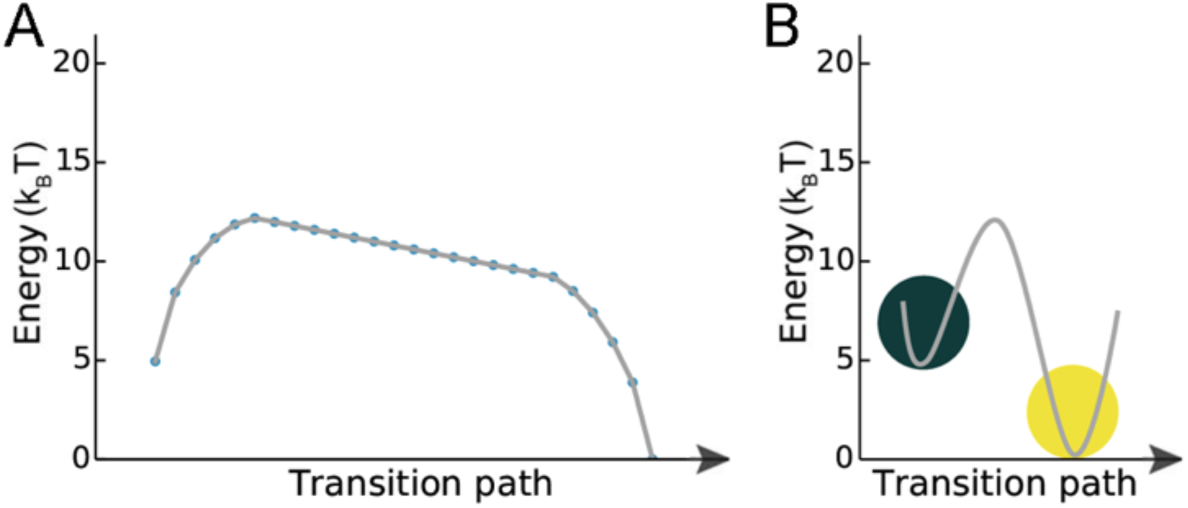
Transition paths from the ground state to the H3K9me3 heterochromatin domain. (A) Transition path constructed using the path deformation approach. The states along the transition path are shown in dots. Details for this calculation are provided in the *Section: Calculating the minimum energy barrier path between chromatin domains using a path deformation approach.* (B) Transition path constructed from the Wang-Landau sampling. This figure is the same as Fig. 5C in the main text.

**Fig. S6.**
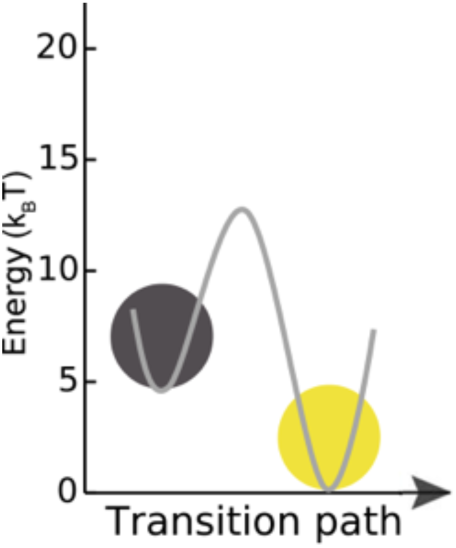
The most probable transition path from the ground state to the H3K27me3 heterochromatin domain. Similar to the path shown in Fig. 5C of the main text for the H3K9me3 heterochromatin, there are no intermediate states along the path, and the transition is a cooperative process.

**Fig. S7.**
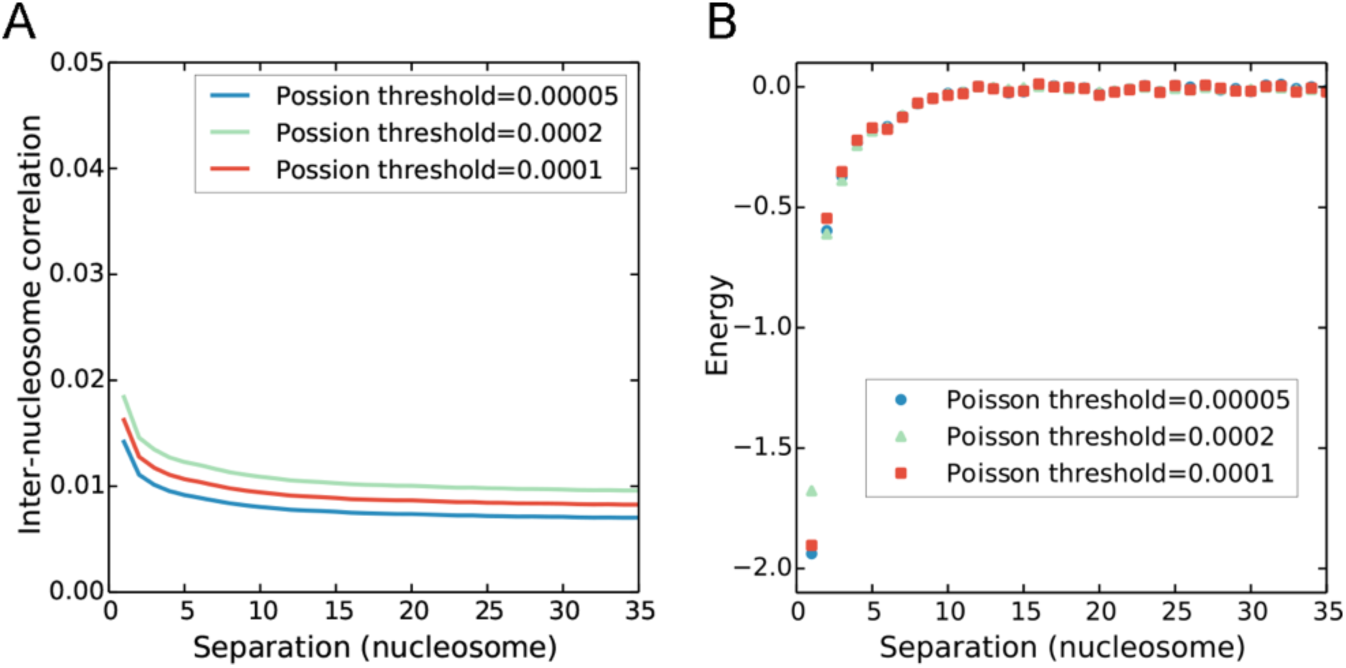
Inter-nucleosome (A) correlation and (B) interaction energies for H3K9me3 as a function of the genomic distance derived from data processed with different Poisson thresholds. See the text *Section: Evaluating the sensitivity of interaction energy strength with respect to the Poisson threshold used in the binarization of experimental data* for details.

